# Evaluation of adipose-derived stromal cell infused modified-hyaluronic acid scaffolds for post cancer breast reconstruction

**DOI:** 10.1101/2025.04.03.646623

**Authors:** Ritihaas Surya Challapalli, Joanne O’Dwyer, Niall McInerney, Michael J Kerin, Garry P Duffy, Eimear B Dolan, Roisin M Dwyer, Aoife J Lowery

## Abstract

**Introduction:** Primary breast cancer surgery can compromise aesthetics and quality-of-life for breast cancer patients. While breast reconstruction improves these outcomes, current methods are limited by suboptimal aesthetic outcomes and potential complication risks. There is an urgent clinical need for improved approaches to post surgical reconstruction for breast cancer patients. Adipose-derived stromal cells (ADSCs) with biological scaffolds are being widely evaluated for tissue engineering applications in the field of reconstruction.

**Aims:** This study aimed to assess the biomechanical properties, biocompatibility, adipogenic potential of ADSCs encapsulated in modified hyaluronic acid derivatives in vitro; and efficacy and tissue integration of this construct in vivo in a murine breast cancer and reconstruction model.

**Methods:** ADSCs were obtained, with informed consent, from female breast cancer patients undergoing autologous breast reconstruction or cosmetic procedures (n=8) aged 47±12 years. Modified hyaluronic acid solution was combined with 1×10^6^ ADSCs/mL and crosslinked using hydrogen peroxide and horseradish peroxidase. Young’s modulus, cell viability and adipogenic potential of the cell-loaded hydrogels were assessed in vitro. In vivo, hydrogels combined with murine ADSCs were grafted into a murine breast cancer model and tissues were harvested for immunohistochemistry after 4 weeks.

**Results:** ADSCs were characterised via morphology, Colony forming unit-fibroblast (CFU-F) assay, flow cytometry and multilineage differentiation. The cell-loaded hydrogels had a compressive Young’s modulus of 7.35±0.96 kPa after 21 days in culture, similar to human breast adipose tissue (∼10 kPa). High ADSC viability was observed after 21 days in culture, and ADSCs differentiated into mature adipocytes. After 4 weeks in vivo, hydrogels exhibited adipocytes, vascular endothelium, and pericyte-like cells.

**Conclusion:** This study demonstrates the potential suitability of modified hyaluronic acid hydrogels encapsulating ADSCs for adipose tissue engineering for post breast cancer reconstruction.

## INTRODUCTION

Surgical tumour excision remains the cornerstone of curative breast cancer treatment, with mastectomy and breast conserving surgery (BCS) the established surgical approaches. Both procedures can result in physical defects which impair the aesthetic appearance of the breasts and may cause altered body image and psychological distress for patients [1]. Breast reconstruction, whether immediate or delayed after mastectomy, as well as volume replacement following BCS, have been shown to enhance psychosocial well-being and improve quality of life outcomes [1–3]. Currently, commonly used breast reconstruction methods including implant-based reconstruction or autologous reconstruction with tissue transfer are limited by potential morbidity and suboptimal aesthetic outcomes [4–9]. Recent advancements in tissue engineering, particularly in the field of adipose tissue engineering, have demonstrated considerable potential for enhancing breast reconstruction procedures. Adipose tissue engineering involves the use of autologous stem/stromal cells combined with biocompatible scaffolds to create an environment that facilitates de novo adipose tissue regeneration [10, 11]. Adipose derived stromal cells (ADSCs) can be effectively combined with biomaterials such as hyaluronic acid hydrogels to generate mature adipose tissue. These hydrogels have the potential to serve as scaffolds, providing mechanical support and creating an environment that promotes optimal cell adhesion, growth, proliferation, and differentiation of ADSCs. [12–14]. An ideal scaffold for adipose tissue engineering should possess biocompatibility, non-immunogenicity, and biodegradability. Furthermore, the scaffold should typically fall within the range of 1-10 kPa, which is necessary for promoting adipogenic lineage commitment in ADSCs [15–17].

Hyaluronic acid (HA) is a non-sulphated glycosaminoglycan composed of alternating units of D-glucuronic acid and N-acetylglucosamine, ubiquitous in the extracellular matrix. HA is a versatile extracellular matrix (ECM) component widely used in biomedical applications due to its role in tissue hydrodynamics, cell proliferation, cell-cell communication, and non-immunogenicity [18]. HA is an ideal biomaterial for regenerative approaches as it can be chemically modified to modulate degradation and the addition of bioactive molecules has been shown to support cell attachment, proliferation and differentiation [17, 19]. Previous studies have shown that chemically modified HA derivatives have the potential to repair cardiac defects [20], cartilage defects [21], central nervous system (CNS) disorders [22] and aid wound repair [23].

The tuneable properties of HA cross-linked using hydrogen peroxide (H2O2) and horseradish peroxidase (HRP) are an attractive option for in situ cross-linkable hydrogels, well-suited for minimally invasive procedures [24–27], particularly in cases such as irregular and variable volume deformities which may be caused by breast cancer surgery. The injectable nature of in situ cross-linkable hydrogels allow for precise placement and adaptation to the specific contours and shape of the breast, with the potential to enhance aesthetic outcomes [28]. Gallagher et al. investigated H2O2/HRP cross-linked HA hydrogels encapsulating bone marrow derived human MSCs in vitro. Their study found that 2% w/v cross-linked HA hydrogels showed improved cell attachment, survival, and angiogenic potential [19]. Dolan et al. further investigated H2O2/HRP cross-linked HA hydrogels encapsulating human ADSCs and demonstrated viability of the cells in vitro and successfully delivered these hydrogels into porcine myocardium via a custom catheter [29]. In another publication by Dolan et al, similar human ADSC loaded HA hydrogels were delivered to the epicardial surface of porcine hearts using a custom bioresorbable patch and found improvements in heart function [20]. These studies suggest that cross-linked HA hydrogels are promising scaffolds for tissue engineering due to their biocompatibility, functional properties and deliverability, however they have never been investigated for adipose tissue engineering.

Building on this previous research, the current study aims to investigate the potential of modified HA cross-linked hydrogels in combination with ADSCs for adipose tissue engineering. We hypothesise that this hydrogel formulation can be utilized to encapsulate and support differentiation of autologous ADSCs for adipose tissue engineering in the context of post cancer breast reconstruction (Figure 1).

**Figure 1.**
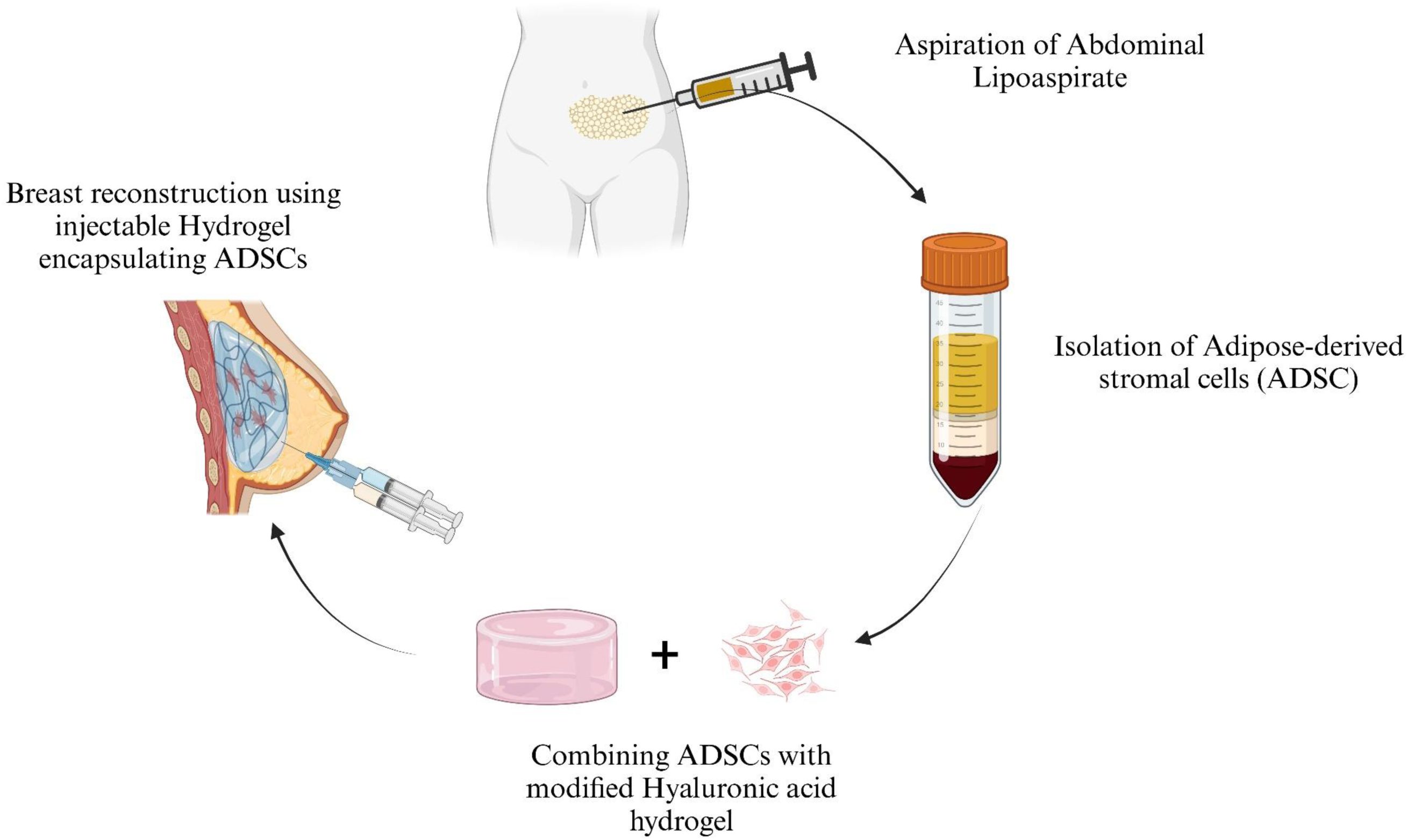
Overview of the clinical application of injectable hydrogel encapsulating adipose-derived stromal cells (ADSCs) for breast reconstruction.

In this study, we assessed the feasibility and efficacy of our approach via in vitro and in vivo adipose tissue regeneration techniques. In vitro, we optimised our human ADSC-loaded-hydrogels and examined mechanical properties, viability, and adipogenic differentiation. Building upon the promising in vitro results, we then assessed the efficacy of murine ADSC-loaded-hydrogels to generate mature fat in vivo using a murine breast cancer excision and reconstruction model. The goal of this approach was to enable evaluation of the tissue reconstruction efficacy of the ADSC-loaded-hydrogel.

## MATERIALS AND METHODS

### Isolation and culture of human ADSCs

Abdominal fat or lipoaspirate was obtained from female breast cancer patients undergoing autologous breast reconstruction or cosmetic procedures (n=8) aged 47±12 years, with informed consent according to the institutional (Galway University Hospital) ethics approval. Fat was minced using surgical blades prior to digestion. Lipoaspirate or minced tissue was washed with PBS, then digested with ∼100 units/mL Collagenase III (Worthington Biochemical Corp. NJ, USA) made up in High glucose DMEM with Glutamax™ I (Gibco) supplemented with 10% v/v Fetal Bovine serum (FBS) (Gibco) and 1% antibiotic-antimycotic solution (Gibco) (Complete media) at 37°C for 1 hr with mechanical agitation using a rotating mixer. This was followed by centrifugation at 1200 rpm for 5 mins and washed with PBS, the pelleted stromal vascular fraction was obtained and cultured in complete media at 37°C, 5% CO2. The culture media was replaced every 2-3 days and the cells were sub-cultured into 1:3 at ∼80% confluency.

### Preparation of hyaluronic acid hydrogel mixture

Hyaluronic acid-hydroxyphenyl-arginylglycylaspartic acid (HA-PH-RGD) and Hyaluronic acid-tyramine (HA-TA) mixed at a ratio of 1:1 as previously described [19] was used in this study, herein referred to as modified-HA. The lyophilised modified-HA obtained from Contipro (Czech Republic), was rehydrated using PBS overnight to produce a 2% w/v final dispersion (this was adjusted based on the volume of cells to be added) and filtered through 0.22 µm filter to sterilise as previously described [19]. ADSCs cultured to passage 3 were then resuspended in an appropriate volume of complete media (to produce the final 2% w/v hydrogel) and mixed with the modified-HA dispersion.

The modified-HA dispersion was divided into two equal parts (A, B); Part A was combined with ADSC suspension and horseradish peroxidase (0.36 U/mL, HRP dissolved in 0.1% bovine serum albumin in PBS). Part B was combined with PBS equivalent to the volume ADSC suspension and 0.1% w/v hydrogen peroxide (H2O2) in PBS. Parts A and B were loaded into luer-lock type syringes [19], connected to a static mixer and depressed parallel into custom pre-fabricated polytetrafluoroethylene (PTFE) moulds, each containing 200 µL. The formulation was polymerised in the mould instantly and the polymerised “ADSC-loaded-hydrogel” were then transferred into a 24-well plate with 1 mL complete media. The “ADSC-loaded-hydrogel” were cultured at 37°C, 5% CO2 and media was changed every 2-3 days. Blank hydrogels (for baseline) were prepared as previously mentioned without the cells i.e., complete media was added to part A of the hydrogel mixture instead of cell suspension.

### Adipogenic differentiation of human ADSCs in the modified HA hydrogels

Encapsulated human “ADSC-loaded-hydrogels” (n=5) and blank hydrogels (n=5) (hydrogel alone without cells) were subjected to adipogenic differentiation (Differentiated) by addition of adipogenic differentiation media containing 1000 nM Dexamethasone, 200 μM Indomethacin, 500 μM 3-isobutyl-1-methylxanthine (IBMX), and 10 μg/mL Insulin as previously described [30], and control hydrogels (± cells) were cultured in complete media (Undifferentiated) in a 24-well plate at 37°C, 5% CO2.

### Evaluation of compressive Young’s modulus

Firstly, the diameter and height of hydrogels from the “Differentiated” and “Undifferentiated” groups ± cells were assessed at day 1, 7 and 21 (n=4/group) using Digital Vernier callipers. Hydrogels were then mechanically tested in unconfined compression between impermeable platens using a standard materials testing machine with a 10 N load cell (Zwick Z005, Roell, Germany) as previously described [19]. Briefly, the hydrogels were transferred to a petri-dish containing complete media and placed on the bottom plate of the mechanical tester. A preload of 0.05N was applied to ensure the contact of testing plates with the hydrogel before recording. The hydrogels were compressed using a strain rate of 10 mm/min and the displacement data was recorded at 10 Hz using testXpertII software. The hydrogels were subjected to a maximum of 30% strain before the test was stopped. The compressive modulus was determined as the slope of the linear elastic region of the resulting stress-strain curve, between 0% and 10% strain.

### Viability and lipid deposition of human ADSCs in the modified HA hydrogels

Cell viability and lipid deposition was assessed in “hADSC-loaded-hydrogel” from the “Differentiated” and “Undifferentiated” groups at day 1, 7 and 21 of the adipogenic differentiation cycle. Each cylindrical hydrogel from both conditions were bi-sected (in transverse plane) and transferred to µ-Slide 8 Well chambered cover slip (ibidi). One section of the hydrogel was incubated with Nile Red Staining solution (Abcam) containing 5 µM Hoechst 33342 (Abcam) for 30 min at RT in the dark. The other section was incubated with Live/Dead staining kit (Invitrogen) containing 4 µM Calcein-AM and 8 µM Ethidium homodimer-1 for 30 min at RT in the dark.

Images of fluorescently labelled cells were acquired with Andor Spinning Disc confocal microscope using IQ3 software. Each hydrogel image is comprised of 201 image slices, 5 µm apart (step size) resulting in a Z-stack of 1 mm high hydrogel. Maximum intensity projections of each z-stack was curated using IQ3 software. ROI of the images were processed using Fiji (ImageJ v1.53c, Bethesda, USA).

### Adipokine secretion

Adipokines are bioactive molecules secreted by adipose tissue that play significant roles in regulating metabolism, inflammation, and overall energy homeostasis [31, 32]. In this study, adipokines secreted by the hADSCs encapsulated in hydrogels were analysed in the conditioned media. Conditioned media (CM) from “hADSC-loaded-hydrogel” (n=3 hydrogels per group) were pooled at Day 1 and 21 of the adipogenic differentiation cycle respectively, centrifuged at 1000 rpm for 1 min and stored at −20°C. Adipokines secreted by the hADSCs encapsulated in hydrogels were analysed in the CM using Proteome Profiler Human Adipokine Array Kit (ARY024, R&D systems) according to manufacturer’s instructions. The membranes were exposed for 240 sec and images were acquired on Bio-rad GelDoc XR+ system using ImageLab software. Densitometry analysis was performed using Fiji (ImageJ, Bethesda, USA) with background correction.

### Isolation, culture and characterisation of murine ADSCs (mADSCs) for in vivo studies

The in-vivo study was approved by the Animal Ethics Committee (University of Galway) and Project and Individual Authorizations were granted by the Health Products Regulatory Authority (HPRA) of Ireland (project license no AE19125/P119).

To obtain a murine ADSC cell population for use in the in vivo model, subcutaneous mammary fat pad was excised from Female Balb/C AnNRj mice (Janvier Labs, France), minced with surgical blades, digested using collagenase III and cultured as previously described [30]. The mADSCs were characterised via morphological analysis to assess their fibroblast-like shape and plastic adherent properties, ability to form colonies via CFU-F assay and tri-lineage differentiation assays (osteocytes, chondrocytes and adipocytes) as previously described [30].

### Murine breast tumour induction, excision, and reconstruction

Female Balb/C AnNRj mice (Janvier Labs, France) aged 6-8 weeks old (n=32), received orthotopic injection of 1×10^5^ 4T1-Luc-2 cells (ATCC) in basal RMPI-1640 into the 4^th^ inguinal mammary fat pad under inhalation anaesthesia using 25G needles and 1 mL syringes. Animals were monitored until a palpable tumour developed (Figure 1).

Once palpable, tumour growth was monitored using Vernier callipers until they reached the appropriate size of 100 ± 50 mm^3^, then animals were randomly assigned to four groups with tumour resection with or without implantation of varying hydrogel/cell combinations (described in Table 1). Primary tumour resection was performed under inhalation anaesthesia. Following tumour removal, the wound was closed using wound clips in resection only group. In the hydrogel-containing groups, pre-formed hydrogels (∼ 200 µL) were gently placed into the cavity left by the tumour resection, and the wound was closed with wound clips. For the cells only group, 1x 10^6^ mADSC cell suspension in 200 µL basal DMEM media was injected into the adjacent tissue after tumour removal and the wound was closed using wound clips.

**Table 1.** Assignment of animal into various treatment groups.

| **Group** | **n** | **Treatment** |
| --- | --- | --- |
| **Resection only** | **8** | **Tumour resection and primary wound closure only (control)** |
| **mADSC loaded hydrogel** | **8** | **Tumour resection and reconstruction with pre-formed cell-hydrogel complex (1x10<sup>6</sup> mADSC in 200 µL hydrogel)** |
| **mADSC only** | **8** | **Tumour resection and reconstruction with injection of 1x10<sup>6</sup> mADSC suspension (in 200 µL DMEM basal media) into the remaining mammary fat pad** |
| **Hydrogel only** | **8** | **Tumour resection and reconstruction with pre-formed hydrogel only (200 µL hydrogel)** |

The animals were euthanized 4 weeks post-reconstruction via cardiac puncture under inhalation anaesthesia. Tumour tissue, tumour-associated mammary fat pad, reconstructed tissue, contralateral disease-free mammary fat pad, spleen and lungs were harvested for IHC (Figure 1).

### Immunohistochemistry

The harvested tissues were immediately transferred into 4% Paraformaldehyde (PFA) in PBS (pH.6.9) and stored at room temperature until processing and embedding. The harvested tissues and the excised hydrogels were processed in series of alcohols and embedded in Paraffin using Excelsior ES tissue processor (Thermo Scientific) and HistoCore Arcadia H embedding system (Leica).

IHC was performed on 5 µm sections using Ventana benchmark Ultra (Roche diagnostics) autostainer. The sections were then incubated with primary antibodies against Perilipin-1 (1:200, ab3526, Abcam), α-SMA (1:200, ab124964, Abcam), F4/80 (1:150, ab111101, Abcam), CD31 (1:50, ab28364, Abcam), Ki67 (1:200, ab15580, abcam). Following the incubation with primary antibody, the sections were incubated with UltraMap Anti-Rabbit HRP (760-4315, Roche) and developed using ChromoMap DAB kit (Roche). Sections were stained with Hematoxylin and Eosin on bench-top. Images were taken using Olympus VS120-L100-W virtual slide scanner (Olympus) and OlyVia v3.3.

### Data and Statistical analysis

All experiments were performed in triplicate. Data is presented as the mean values ± standard deviation, unless otherwise specified. One-way ANOVA with Tukey’s multiple comparison was performed to analyse the difference between groups and p < 0.05 was considered to be significant. Images were processed using Fiji (Image J) Software, QUPath and statistical analysis was performed using GraphPad Prism v9.3.

## RESULTS

### Isolated ADSC were found to be an appropriate cell source for the hydrogel construct

All isolated ADSCs were confirmed to be plastic adherent, showed spindle shaped morphology (Figure 2A(i)), formed colonies (Figure 2A(ii)), exhibited multilineage differentiation (Figure 2B (i-iii)) and found to be >95% positive for CD73, CD105 and CD90 and <5% or negative for CD 45 and CD34, using flow cytometry (Figure 2C).

**Figure 2.**
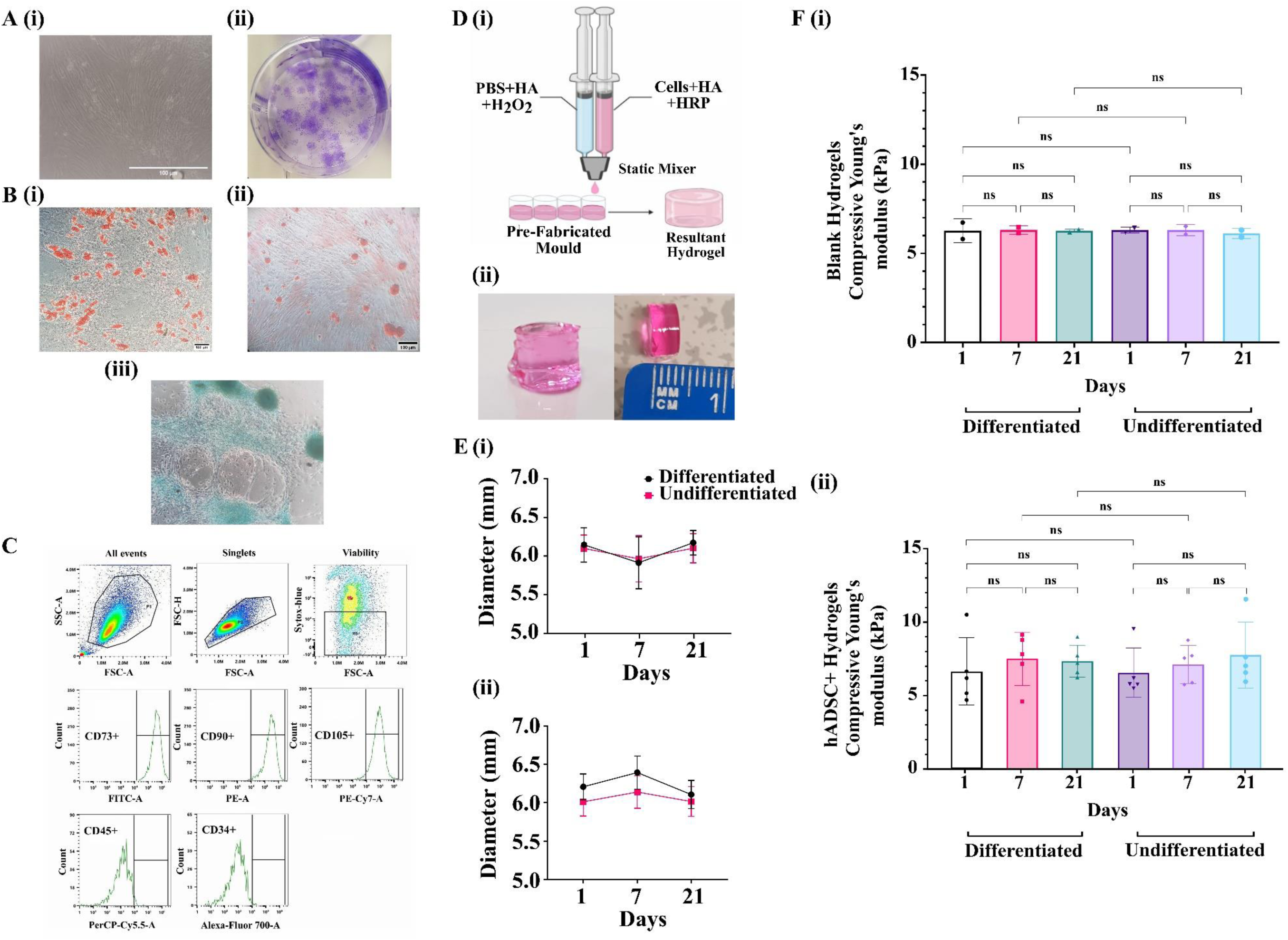
Isolation, characterisation of hADSCs and manufacturing of stable hydrogels (A)Characterisation of hADSC (i)Morphology, (ii) Colony forming unit-fibroblast assay (CFU-F); B(i) Adipogenic differentiation (ii) Osteogenic Differentiation (iii) chondrogenic differentiation, scale 100 μm; (C) Immunophenotyping depicting expression of CD73, CD90, CD 105 and absence of CD 34 and CD45;(D) (i) Schematic representation of preparation of hydrogel to encapsulate ADSCs using benchtop hydrogel mixer (ii) Macroscopic image of hADSC encapsulated hydrogel; (E)(i) Diameter of blank hydrogels subjected to differentiated vs undifferentiated conditions measured at Day 1,7, 21 of adipogenic cycle (n=2); (ii) Diameter of hADSC encapsulated hydrogels subjected to differentiated vs undifferentiated conditions measured at Day 1,7, 21 of adipogenic cycle (n=5); (F)(i) Young’s modulus calculated from 0-10% compressive strain of Blank Hydrogels (n=2) vs (ii) Hydrogels encapsulating hADSCs (n=5) subjected to adipogenic differentiation media (Differentiated) and complete media (Undifferentiated), mean ± SD, kPa-Kilopascal; HA: Hyaluronic acid, PBS: Phosphate-buffered saline, HRP: Horseradish peroxidase, H202: Hydrogen peroxide.

The modified-HA hydrogels displayed instant crosslinking within the mould and maintained a cylindrical shape (Figure 2D (i-ii)). These hydrogels were transferred to a 24-well plate and cultured with complete media. Under an inverted microscope, the ADSCs encapsulated in the hydrogel were clearly visible.

### Compression modulus of modified hyaluronic acid hydrogels was similar to native breast adipose tissue (∼10 kPa)

The measured diameter of blank hydrogels (without cells) exposed to adipogenic differentiation media remained stable over time, with values of 6.20 ± 0.16 mm on Day 1, 6.39 ± 0.21 mm on Day 7, and 6.10 ± 0.18 mm on Day 21. Similarly, in hydrogels subjected to complete (undifferentiated) media, diameters were consistent across time points (ranging between 6.01 and 6.13 mm). No significant differences in diameter were observed between differentiated and undifferentiated groups at any time point. (Figure 2E (i)).

For hADSC-loaded hydrogels, a similar trend was observed. When exposed to differentiation media, diameters show minimal change from 6.14 ± 0.22 mm on Day 1 to 5.91 ± 0.34 mm on Day 7, before stabilizing at 6.17 ± 0.16 mm by Day 21. Hydrogels in the undifferentiated group also showed minimal change, ranging from 6.10 ± 0.17 mm to 5.97 ± 0.30 mm from Day 1 to 21. Importantly, no significant differences in diameter were observed between differentiated and undifferentiated groups at any time point (Figure 2E (ii)).

The compressive Young’s modulus (0-10% strain) of blank hydrogels in both differentiation and undifferentiated media remained consistent over the study period, with values ranging between 6.10 and 6.30 kPa across all time points and no significant differences were observed (Figure 2F (i)).

For hADSC-loaded hydrogels, a slight increase in stiffness was observed in both groups over time. In the differentiated group, the modulus increased from 6.56 ± 2.05 kPa on Day 1 to 7.35 ± 0.96 kPa by Day 21. The undifferentiated group followed a similar trend, with values ranging from 6.55 ± 1.50 kPa on Day 1 to 7.71 ± 0.99 kPa on Day 21. No statistically significant differences were detected between the differentiated and undifferentiated groups at any time point (see Figure 2F (ii)).

### hADSCs encapsulated in the hydrogels were viable and differentiated into adipocyte lineage at day 21

At Day 1, 7 and 21, the hADSCs encapsulated in hydrogels were found to be viable, as indicated by the presence of green-fluorescent cells (labelled with Calcein-AM). In contrast, dead cells were visualized as red-fluorescent cells (labelled with ethidium homodimer-1). This observation was consistent in both the differentiated and undifferentiated group (Figure 3A).

**Figure 3.**
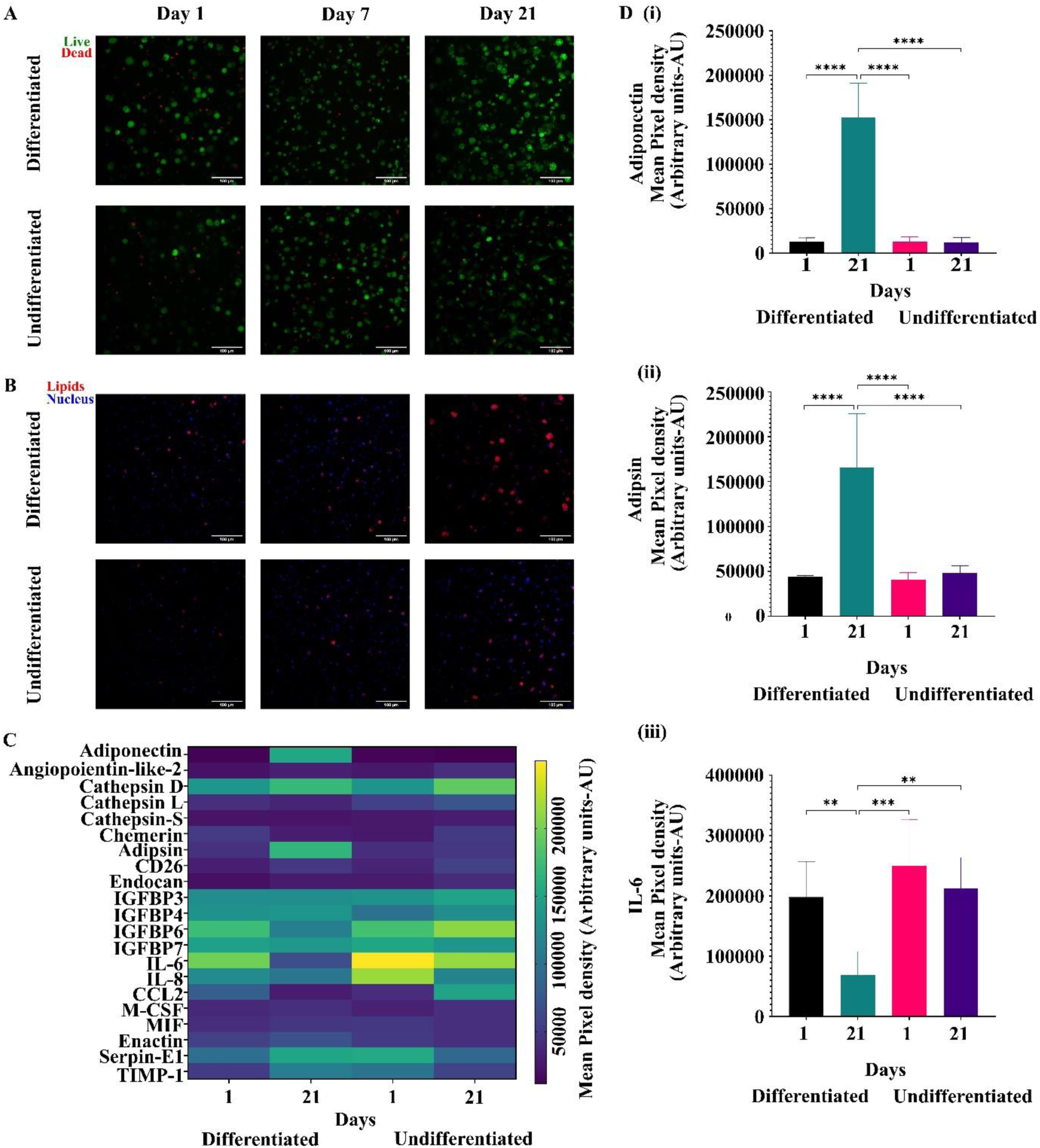
Viability and adipogenic differentiation of hADSCs encapsulated in the hydrogel over 21 days. (A) Representative max intensity projection of confocal images of hydrogels showing live cells (green) and dead cells (red). Scale 100 µm; (B) Representative max intensity projection of confocal images of hydrogels showing nucleus (blue) and lipids (red). Scale 100 µm;(C) Heatmap of Mean pixel density ± SD of Adipokine secretions from Cell-hydrogel conditioned media subjected to Adipogenic differentiation (Differentiation) or complete media (undifferentiated)at Day 1 and Day 21; (D) (i) Mean pixel density ± SD of Adiponectin secreted in cell-hydrogel conditioned media across groups; (ii) Mean pixel density ± SD of Adipsin secreted in cell-hydrogel conditioned media across groups; (iii) Mean pixel density ± SD of IL-6 secreted in cell-hydrogel conditioned media across groups, mean pixel density ± SD in Arbitrary units corrected for background. ** p<0.05, *** p< 0.001, **** p<0.0001.

No necrotic regions with clustered dead or dying cells were detected at any time point in differentiated or undifferentiated hydrogels (Figures 3A).

At Day 1 the differentiated hydrogels did not exhibit any noticeable lipid accumulation but started showing indistinct lipid droplets. At Day 21, the differentiated hydrogels displayed noticeable accumulation of lipid droplets, observed as distinct spherical clustered droplets surrounding the nucleus (Figure 3B, Day 21 differentiated). In contrast, the undifferentiated hydrogels did not exhibit any significant accumulation of lipid droplets at any time point.

### Conditioned media from hADSC-loaded-hydrogel cultured in differentiation media showed significant increase in levels of secreted adipokines

The conditioned media from hADSC-loaded-hydrogels was analysed for 21 adipokines in both the differentiated and undifferentiated groups at Day 1 and Day 21, as depicted in Figure 3C. The results showed that the conditioned media from the differentiated hydrogels exhibited a significant increase in the mean pixel density (arbitrary units - AU) of Adiponectin (ANOVA with post hoc Tukey test: p<0.0001, Figure 3D (i)) and Adipsin (p<0.0001, Figure 3D (ii)) at Day 21 compared to the other groups.

Conversely, there was a significant decrease in the mean pixel density (AU) of IL-6 at Day 21 of the differentiated hydrogels compared to Day 1 of differentiated (p=0.0051), Day 1 of undifferentiated (p=0.0002), and Day 21 of undifferentiated (p=0.0020) (Figure 3D (iii)).

### Weight and volume of reconstructed tissue was significantly increased when mADSC-loaded-hydrogels were used compared to the mADSCs alone

4 weeks post tumour excision and reconstruction procedure, the reconstructed tissues were excised for analysis (n=31, 1 excluded due to humane endpoint) (Figure 4A).

**Figure 4.**
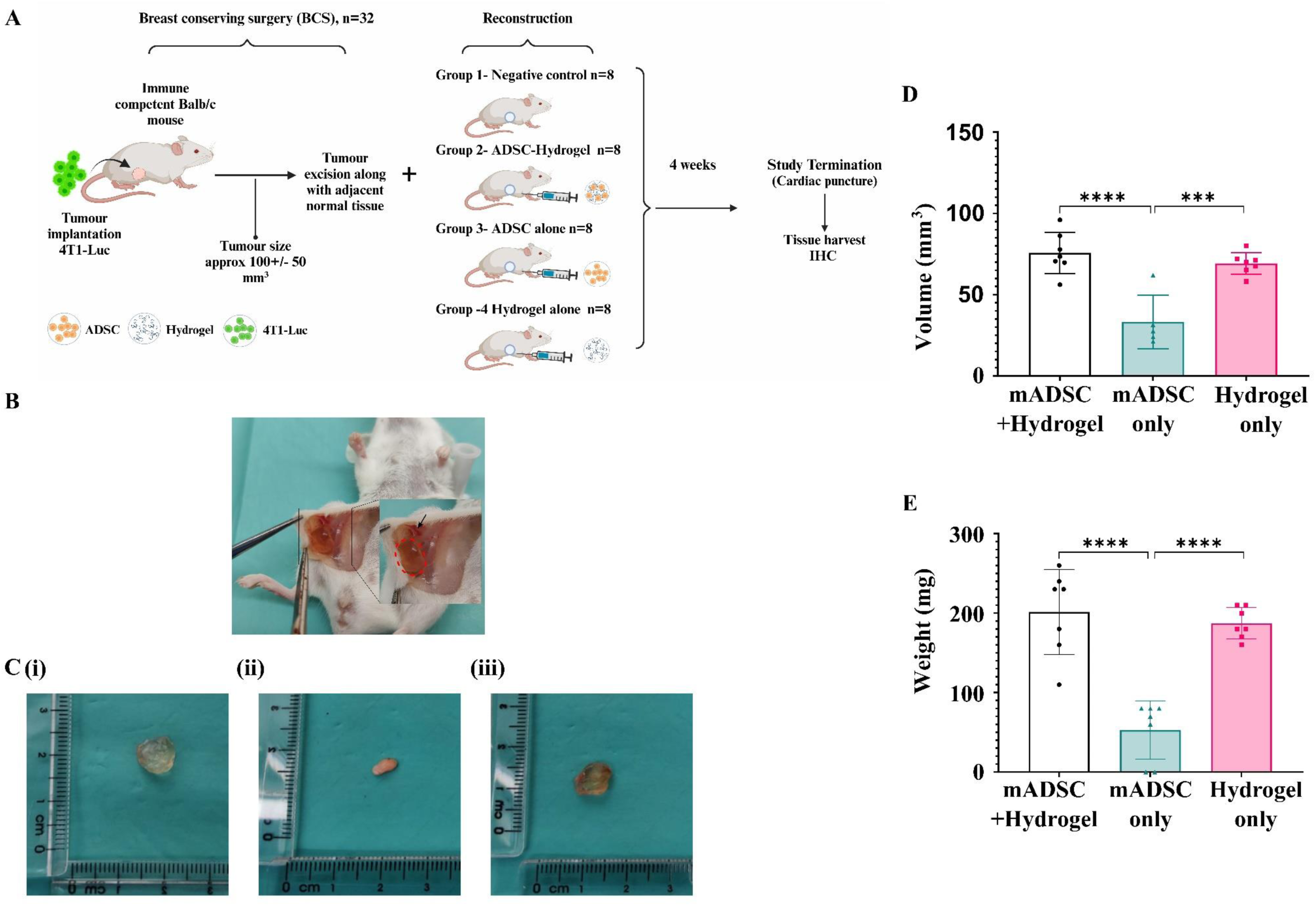
Assessment of invivo biocompatibility and integration of hydrogels encapsulating mADSCs (A) Immuno-competent Balb/C mice were subcutaneously implanted with 4T1-luc cells to induce tumour followed by resection of tumour and reconstruction with hydrogel encapsulating mADSCs, mADSCs only and hydrogel only compared to resection only control for 4weeks and tissue harvest; (B) Representation of the Hydrogel location in the mice (red dotted line in the inset) held by connective tissue (black arrow in the inset); (C) Gross necropsy images of excised reconstructed tissue (i) hydrogels encapsulating mADSC, (ii) tumour adjacent fat pad that received mADSC as suspension (iii) hydrogel only (without cells); (D) volume (mm^3^) of excised reconstructed tissue from Group 2 (mADSCs+hydrogel) (n=7), Group 3 (mADSC only) (n=5) and Group 4 (hydrogel only) (n=7); (E) Weights (mg) of excised reconstructed tissue from Group 2(mADSCs+hydrogel) (n=7), Group 3 (mADSC only) (n=5) and Group 4 (hydrogel only) (n=7); Mean ± SD, **** p<0.0001, *** p=0.0001.

The location of the hydrogel was visually confirmed, as shown in Figure 4(B). The polymerised hydrogel remained in the cavity left by tumour removal, and integrated in the host (held in place) by surrounding connective tissue, as indicated by the red dotted line and black arrow, respectively (Figure 4B). Gross necropsy images in Figure 4C(i-iii) show excised reconstructed tissues, highlighting differences in reconstruction volume between groups immediately after excision. In the resection only group (negative control), no tissue/fat regrowth in the fat pad was observed in any animal (n=8). Regrowth was observed in all other groups. Two excised hydrogel samples were excluded from the final analysis: one from the “mADSC-loaded-hydrogel” group due to loss of the hydrogel following disruption of wound clips and another from the “hydrogel-only” group lost during the excision process. Data from the remaining samples were included for analysis.

The mean volume of excised reconstructed tissues from “mADSC-loaded-hydrogel” group was 75.68 ± 11.8 mm^3^ (n=7), from “hydrogel-only” group was 69.18± 6.22 mm^3^ (n=7), and from “mADSC-only” group was 33.24± 14.78 mm^3^ (n=5). The volume of reconstructed/regenerated tissue was significantly higher in “mADSC-loaded-hydrogel” group and “hydrogel-only” group compared to “mADSC-only” group (ANOVA with post hoc Tukey test: mADSC loaded hydrogel group vs. “mADSC-only” group p<0.0001, “hydrogel-only” group vs. “mADSC-only” group p=0.0003). No significant difference was observed in the volume of reconstructed/regrown tissue in groups that received the hydrogel ie., between “mADSC-loaded-hydrogel” group and “hydrogel-only” group (ANOVA with post hoc Tukey test: p=0.58) (Figure 4D).

The mean weight of excised reconstructed tissues from “mADSC-loaded-hydrogel” group was 201.43 mg ± 49.4 mg (n=7), from “hydrogel-only” group was 187.14± 18.29 mg (n=7), and from “mADSC-only” group was 53± 34 mg (n=7). Three mice from “mADSC-only” group exhibited no tissue regeneration. The weight of reconstructed/regenerated tissue was significantly higher in “mADSC-loaded-hydrogel” group and “hydrogel-only” group compared to “mADSC-only” group (“mADSC-loaded-hydrogel” group vs. “mADSC-only” group p<0.0001, “hydrogel-only” group vs. “mADSCs-only” group p=0.0001). No significant difference was observed in the weight of reconstructed/regenerated tissue in groups that received the hydrogel ie., between “mADSC-loaded-hydrogel” group and “hydrogel-only” group (ANOVA with post hoc Tukey test: p=0.73) (Figure 4E).

### Capsule formation was observed around all excised hydrogels

H&E and α-SMA staining of the excised hydrogel sections showed the formation of fibrotic capsules around each hydrogel. The thickness of the capsules was measured at five random locations on each section stained with H&E (Figure 5A, B). The mean capsule thickness for “mADSC-loaded-hydrogel” group was found to be 121.6 ±69 µm and the mean capsule thickness for “hydrogel-only” group was 131.2±80 µm. There was no significant difference in capsule thickness between the groups (t-test: p=0.673) (Figure 5C).

**Figure 5.**
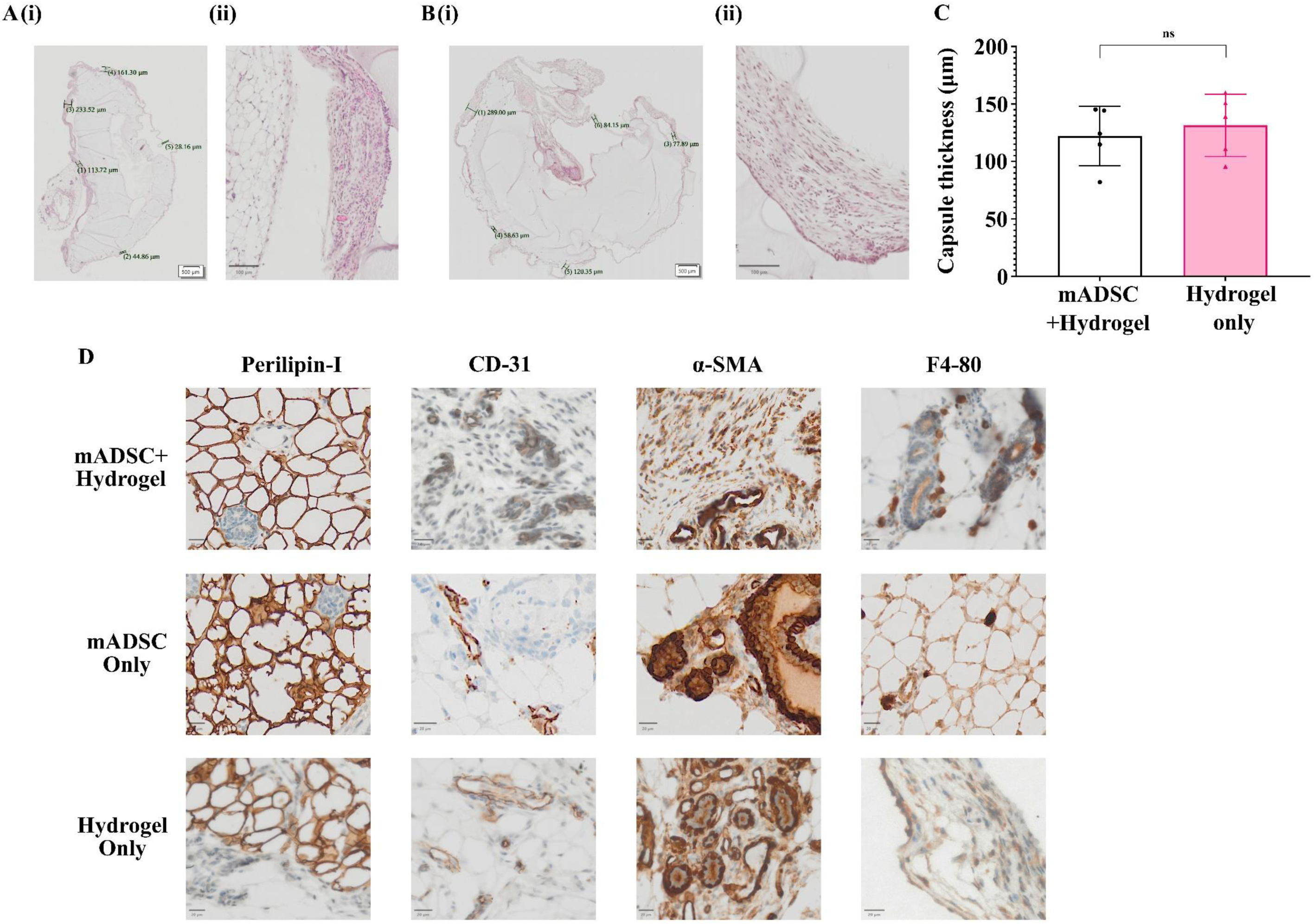
Assessment of biocompatibility of the hydrogel in-vivo. (A) (i) Micrograph of mADSC+hydrogel stained with H&E showing capsule thickness measurements at 5 random locations, Scale 500 µm.; (ii) Magnified image of the capsule, scale 100 µm; (B) (i)Micrograph of hydrogel only stained with H&E showing capsule thickness measurements at 5 random locations, Scale 500 µm.; (ii) Magnified image of the capsule, scale 100 µm; C) capsule thickness compared between mADSC+Hydrogel and Hydrogel only groups. (D) Immunohistochemistry staining of hydrogels in mADSC+Hydrogel, mADSC only and hydrogel only groups stained against Perilipin-1 showing adipocytes; CD31 staining microvascular structures; α-SMA staining myofibroblasts and myoepithelial like-cells focussed around vascular and ductal structures; F4/80 staining infiltrating macrophages. Scale 20 µm.

### Excised hydrogels exhibited mature adipose tissue and formation of microvascular structures

Immunohistochemistry revealed formation of mature adipocytes in both “mADSC-loaded-hydrogel” group and “hydrogel-only” group visualised by staining against Perilipin-1 (Figure 5D). CD31 staining showed formation of microvascular structures with distinct lumen in both “mADSC-loaded-hydrogel” group and “hydrogel-only” group (Figure 5D). α-SMA staining revealed myofibroblasts in the capsules and myoepithelial-like cells surrounding luminal structures in both groups (Figure 5D). F4-80 staining showed infiltration of macrophages in both “mADSC-loaded-hydrogel” group and “hydrogel-only” group (Figure 5D).

IHC of excised tissue from “mADSC-only” group also revealed presence of mature adipocytes, CD31+ microvascular structures, α-SMA+ myoepithelial like cells and F4-80+ macrophages (Figure 5D). However, it should be noted that the reconstruction procedure was similar to breast conserving surgery, where the tumour was excised from the adjacent fat tissue and mADSCs in suspension were injected into existing native fat tissue. So, it is difficult to differentiate between native mature fat vs regeneration of tissue from the injected mADSCs.

## DISCUSSION

In this study we have assessed the suitability of a modified hyaluronic acid hydrogels formulation to encapsulate and support differentiation of autologous ADSCs for adipose tissue engineering in the context of post cancer breast reconstruction.

This work demonstrates that the HA hydrogel is a suitable scaffold to support ADSC differentiation to mature fat in vitro and in vivo. The hydrogels which encapsulated ADSCs isolated and cultured in complete media until passage 3 resulted in cylindrical hydrogels that were optimally cross-linked. These hydrogels were easily separable from the mould and maintained their geometric shape throughout the duration of the study. Throughout the 21-day in vitro study, the diameter of the HA hydrogels +/- ADSCs remained stable in both differentiated and undifferentiated groups.

The absence of significant swelling or degradation in the hydrogel is paramount as it ensures safe interactions with biological tissues, it also confirms stability, which is crucial for medical applications and tissue engineering where materials must endure for extended periods, facilitating successful and predictable outcomes. Furthermore, the compressive Young’s modulus of the ADSC-loaded hydrogels consistently ranged between 6-8 kPa. This range is considered ideal for facilitating the commitment of ADSCs to the adipocyte lineage [36]. Alkhouli et al evaluated compression modulus of excised subcutaneous adipose tissue from 25 female patients undergoing elective surgery with a mean age of 48.3 ± 12.2 yrs. The study reported an elastic modulus of 11.7±6.4 kPa when up to 30% strain was applied using a custom built uniaxial tensile tester [37]. Umemoto et al. evaluated non-linearity of elastic properties of breast tissue and reported an elastic modulus of 9.25± 2.56 kPa when 0.6-0.8 kPa stress was applied using Instron 3342 tensile tester with a 10-N load cell and φ3 mm cylindrical indenter [38]. In this study we have demonstrated that the mechanical properties of these hydrogels are within the range of native tissue and suitable to provide an appropriate biomechanical environment necessary for ADSCs to commit to adipogenic lineage, when supplemented with adipogenic differentiation media.

The live/dead staining analysis revealed that the cells remained viable throughout the 21-day in vitro study, with a visibly low number of dead cells observed. Importantly, no necrotic regions with clustered dead or dying cells were detected at any time point. Gwon et al. conducted a study where ADSCs were encapsulated in Heparin-Hyaluronic acid hydrogels. They reported that under undifferentiation conditions, the cells remained viable, exhibited proliferation, and were evenly distributed throughout the hydrogel at Day 1, 7, 14 and 21, as visualized through Live/Dead staining [39]. Our findings align with these and suggest that our modified HA hydrogel formulation provides a suitable environment to support the supply of nutrients and gases required for the viability of the cells throughout the culture period, and survival of the graft.

The Nile Red staining analysis confirmed a significant increase in lipid deposition in the cells undergoing differentiation compared to the control group in our 21-day in vitro study. This observation was supported by the presence of micro-clustered lipid droplets within each cell, with the nucleus located in the periphery in differentiated ADSC-loaded-hydrogel at day 21, resembling the characteristic morphology of adipocytes. Additionally, the significant increase in the secretion of Adipsin and Adiponectin observed in the conditioned media from the “hADSC-loaded-hydrogel” in the differentiation group at day 21 further confirms the successful differentiation of hADSCs into mature adipocytes. Adipsin and Adiponectin are involved in regulating adipose tissue homeostasis. Adipsin is part of the complement system, influencing storage of triglycerides in adipocytes by enhancing glucose transport into the cells and inhibiting lipolysis [33]. Adiponectin is essential for modulating glucose levels and fatty acid breakdown. Monitoring these adipokines provides insights into adipogenesis and the functional capacity of hADSCs [34, 35]. Interestingly, there was a significant decrease in IL-6 at day-21 of differentiation when compared to control and day 1 (both conditions). These results are in line with the literature that suggests ADSCs/MSCs secrete high levels of IL-6 to maintain their regenerative potential and skew differentiation [40]. A study by Kondo et al demonstrated that high amounts of IL-6 was secreted into the culture supernatant at the initiation of chondrogenic differentiation but significantly declined after terminal differentiation [40]. Our study shows high levels of IL-6 at day 1 of the adipogenic differentiation cycle in both differentiated and undifferentiated hydrogels, where the cells have maintained their ADSC-phenotype but the levels significantly decreased after terminal differentiation to adipocyte lineage. These findings are also supported by high levels of IL-6 in undifferentiated ADSC-hydrogels at day 21, and combined with the increased secretion of adipsin and adiponectin provides further reassurance regarding the functionality of the differentiated cells in ADSC-hydrogels at the end of differentiation cycle.

Building on our positive in vitro findings, the objective of the in vivo study was to assess the feasibility and efficacy of modified hyaluronic acid hydrogels encapsulating mADSCs as a post cancer breast reconstruction approach in a murine model of breast cancer. The weight and volume of excised reconstructed tissues was found to be significantly higher in all groups with hydrogel compared to ADSCs alone, indicating the successful role of the hydrogel in providing mechanical architecture for tissue assembly and potential in facilitating volume retention. While the incorporation of hydrogel inevitably contributes to an increase in the weight and volume of the reconstructed tissue, it is also important to emphasize that the hydrogel demonstrated notable resistance to degradation and resorption, 4 weeks post-reconstruction as visually evidenced by the excised tissue. This observation has significant implications in the context of clinical translation, as conventional reconstruction techniques, such as fat grafting or ADSC-enhanced fat grafting, often face limitations related to graft resorption. The introduction of hydrogel presents a promising approach for addressing this issue and enhancing the durability of tissue reconstruction. However, the H&E staining 4 weeks post-reconstruction, revealed a sparse population of cells in both hydrogel groups, primarily concentrated at the periphery of the hydrogel. This observation suggests that during the initial phase of reconstruction, the cells in “mADSC-loaded-hydrogels” group may have migrated and remained at the periphery of the hydrogel in search of nutrients until the formation of microvascular structures. In the case of the “hydrogel-only” group, the native cells surrounding the hydrogel might have attempted to infiltrate but remained at the periphery due to nutrient supply limitations. This phenomenon is commonly observed in in vivo engraftment of biomaterials for tissue engineering, where cells initially redistribute within the hydrogel to establish an adequate nutrient supply [41–43]. As microvascular structures develop, cells may reorganize and proliferate throughout the hydrogel, contributing to tissue formation and functional outcomes [41–43]. Immunohistochemistry of excised tissues from both hydrogel groups showed the presence of CD31+ endothelium and α-SMA+ pericyte-like cells. This observation substantiates the potential for microvascular and vascular structures crucial for the stable integration of the hydrogel and facilitation of tissue regeneration. However, the limited reconstruction time of 4 weeks may not have been sufficient for the cells to fully infiltrate and populate the hydrogel, for the formation of a denser tissue structure. Extending the study termination point to 8-12 weeks may allow us to observe an increase in cell density within our hydrogels.

The excised hydrogels from both hydrogel groups showed the formation of fibrotic capsules, which is characteristic of the foreign body response [44]. However, the thickness of the capsule was not significantly different between the groups, suggesting that mADSCs did not have a significant impact on capsule formation, at least at the analysed time point. The cellularity within the capsule appears similar between the two groups that received the hydrogel, with no overt visual differences in cell density or organization. Microscopic visualization revealed infiltration of F4/80+ macrophages in the hydrogel, mainly concentrated in and around the capsule. Further studies to compare the rate of capsule formation and the thickness of capsules between hydrogel and conventional implants could facilitate clinical translation. Swartzlander et al conducted a host response study on NIH/3T3 fibroblasts encapsulated in Poly-ethyleneglycol (PEG)-based hydrogel in C57bl/6 mice to evaluate the severity of foreign body response (FBR) and subsequent role in graft survival [44]. The study demonstrated that the encapsulated cells triggered acute inflammatory response leading to proportional increase in recruitment of inflammatory cells that reduced over time [44]. The visual analysis in our study indicated that the F4/80+ macrophages were predominantly concentrated in the capsule surrounding the hydrogel rather than in the core of the hydrogel itself.

As a consequence of the sparsely populated hydrogel matrix, the IHC staining showed dye entrapment, where the DAB stain became trapped in the matrix, posing challenges in conducting quantitative cell population analysis. Perilipin-1 staining revealed the presence of mature adipocytes around the periphery of the excised hydrogels in both groups, whether or not they had a capsule surrounding the mature fat. This suggests that the hydrogel provided support for in vivo adipocyte formation and localization. Moreover, the observation of endothelial cells (CD31+) around the adipocyte cell area suggests the development of microvasculature and vascular structures, indicating a potential nutrient and blood supply to the hydrogel system. The presence of CD31+ cells in both hydrogel-containing groups indicates that the hydrogels possess adequate porosity, facilitating cytokine signalling and endothelial infiltration to support vascular ingrowth. This finding aligns with the literature, where Tokatlian et al. demonstrated that porous hyaluronic acid hydrogels (60 and 100 µm porosity) promoted angiogenesis and accelerated wound closure compared to nonporous hydrogels after two weeks of implantation in a diabetic wound healing mouse model [45].

The α-SMA staining of the excised hydrogel sections showed the presence of myofibroblasts, a characteristic feature of the capsule, along with myoepithelial-like cells surrounding luminal structures. This suggests the formation of contractile ductal structures within the hydrogel. Furthermore, visual comparison of micrographs revealed the presence of α-SMA+ cells around the same location as microvascular endothelial (CD31+) cells. Co-localization of α-SMA+ cells around endothelial cells suggest that these cells could possibly be pericytes which are important for formation of contractile vascular structures [46, 47]. Due to the fragile and sparsely populated structure of the hydrogel sections at 4 weeks post-reconstruction, performing multiplex staining on the same sample was not feasible to directly demonstrate the co-localization of endothelial cells and pericytes.

Together the presence of endothelial cells, pericytes and ductal myoepithelial-like cells closer to the fibrous capsule provides further confirmation that the hydrogels from both groups (excised at 4 weeks) were in the granulation phase of the ‘*temporal sequence of inflammation and wound healing*’ [48]. This observation implies that the acute inflammatory response triggered by the cells in and around the hydrogel had started to resolve, and the tissue was undergoing a transition towards a more stable state. The resolution phase of the FBR is an essential process in tissue regeneration, as it indicates the gradual restoration of tissue homeostasis and reduction in the inflammatory response [48, 49]. This is consistent with the resolution phase of the FBR observed in the hydrogel, further supporting the idea that the tissue was undergoing active tissue remodelling and repair [48, 50]. The involvement of these specific cell types in the granulation phase is crucial for tissue regeneration and vascularization, supporting the notion that the hydrogel was in the process of establishing a stable and functional tissue structure.

## CONCLUSION

In conclusion, this study highlights the potential of modified hyaluronic acid hydrogels encapsulating adipose-derived stem cells (ADSCs) for enhancing adipose tissue engineering in breast cancer patients undergoing post-resection reconstruction. The ADSC-loaded-hydrogels retained their geometric shape, exhibited ideal mechanical properties, and supported adipogenic differentiation with stable cell viability, lipid deposition, and adipokine secretion in vitro.

In vivo, the hydrogels facilitated adipocyte formation and microvascular structures, though cell distribution was limited to the periphery, likely due to the 4-week reconstruction period. While macrophages were present and the foreign body response was progressing, mADSCs did not significantly influence this pathway.

Further long-term studies are required to optimize cell density, prolong hydrogel residence, and ensure complete tissue regeneration. Additionally, larger animal models, such as porcine, should be considered for evaluating large-volume reconstruction and ensuring oncological safety. Taken together the results of this study indicate for the first time the potential that ADSC-loaded-HA hydrogels hold for improving breast reconstruction outcomes.

## Acknowledgements

All flow cytometry experiments were performed in the University of Galway Flow Cytometry Core Facility which is supported by funds from University of Galway, Science Foundation Ireland, the Irish Government’s Programme for Research in Third Level Institutions, Cycle 5 and the European Regional Development Fund.

The authors wish to thank the BRU (Bio-Resources Unit) technical, veterinary and administrative staff in University of Galway for facilitating in-vivo studies and for their ongoing assistance, advice and support in animal procedures husbandry, care and welfare.

The authors acknowledge the facilities and scientific and technical assistance of the Anatomy imaging and Microscopy Facility at the University of Galway (https://imaging.universityofgalway.ie/imaging/)

## Data availability

Raw data that support findings of this study are available from the corresponding author upon reasonable request

**Tables and figures: Evaluation of adipose-derived stromal cell infused modified-hyaluronic acid scaffolds for post cancer breast reconstruction**

